# Protein secretion zones within the Gram-positive cell wall

**DOI:** 10.1101/2022.09.09.507258

**Authors:** Manuel Strach, Felicitas Koch, Svenja Fiedler, Klaus Liebeton, Peter L. Graumann

**Author notes:** for correspondence, ^#^.

## Abstract

Whereas the translocation of proteins across the cell membrane has been thoroughly investigated, it is still unclear how proteins cross the cell wall in Gram positive bacteria, which are widely used for industrial applications. We have studied secretion of α-amylase AmyE within two different *Bacillus* strains. We show that a C-terminal fusion of AmyE with the fluorescent reporter mCherry is secreted via discrete patches with very low dynamics that are visible at many places within the lateral cell wall, for many minutes. Expression from a high copy number plasmid was required to be able to see these structures we term “secretion zones”. Zones corresponded to visualized AmyE activity on the surface of cells, and frequently overlapped with SecA signals, but did not necessarily co-localize with the secretion ATPase. Single particle tracking showed much higher dynamics of SecA and of SecDF, involved in AmyE secretion, at the cell membrane than AmyE. These experiments suggest that SecA initially translocates AmyE molecules through the cell membrane, and then diffuses to a different translocon. Single molecule tracking of SecA show changes in distinct diffusion states of SecA, but suggest that AmyE overexpression does not overwhelm the system. Because secretion zones were only found during the transition to and within stationary phase, diffusion rather than passive transport based on cell wall growth from inside to outside, releases AmyE and thus probably secreted proteins in general. Our findings suggest active transport through the cell membrane and slow, passive transition through the cell wall in Gram positive bacteria.

## Introduction

Members of the genus *Bacillus* are famous for their use in industrial production of exoenzymes, and are widely used in biotechnological applications. Protein secretion is a two-step process, involving transport across the cell membrane, and passage through the several-layered peptidoglycan (PG) cell wall. While the prior is relatively well-understood ^1,2^, the latter path is essentially unclear.

It has been estimated that 10% of the encoded *B. subtilis* proteins are secreted ^3^, to make extracellular polymers available for nutrient uptake, with α-amylase being one with widest economic importance ^4^. While the twin-arginine translocation (Tat) pathway requires transported proteins to be folded ^5^, most proteins are secreted in an unfolded state via the general secretory (Sec) pathway ^6^. Proteins destined for the Sec-pathway have an N-terminal signal peptide that delays folding in the cytoplasm ^7^. The SecY, SecE and SecG proteins together form the translocon complex SecYEG, an hourglass-shaped pore in the cell membrane with a constricted ring in the center ^8^. Another component often described as the ‘motor’ that drives translocation is the ATPase SecA ^9^. SecA can interact with both the pre-protein to be secreted and SecYEG ^10^ as it catalyzes the translocation of the polypeptide chain through ATP binding and hydrolysis ^1^. Additionally, the stabilizing protein SecDF plays an important role maintaining a high capacity of protein secretion ^11,12^. To be released from the membrane the signal peptide of the preprotein has to be removed by a signal peptidase ^13^. The two major signal peptidases recognizing the signal peptide of secreted proteins are SipS and SipT ^14^. Furthermore, there are secretion-assisting factors like the membrane-associated, chaperone-like lipoprotein PrsA ^6^. PrsA is crucial for efficient secretion of a number of exoproteins like amylases ^15^.

After overcoming the membrane, the passage through the cell wall is the next barrier for extracellular proteins. The Gram positive cell wall has been described to form a sieve-like meshwork, which allows diffusion of proteins up to a molecular weight of 25 kDa ^16^. However, secreted enzymes have a range of sizes between 15 to 70 kDa, such as amylases, lipases and proteases, so their passage through multiple layers of peptidoglycan would require a pore-forming transport system, or otherwise heterogeneous meshwork-sizes to allow for diffusion passages. Indeed, large cavities within the cell wall, and heterogeneous density of PG strands have been visualized using atomic force microscopy ^17,18^. Such discontinuities within the PG would be compatible with passages for larger proteins through the cell envelope. However, the passage of proteins through the cell wall is still an unresolved question.

The cell wall protects the cell against environmental stress, from bursting due to internal turgor pressure and is responsible for cell shape ^19^. The Gram-positive cell wall consists of several layers and is 30 - 100 nm thick, 30 – 40 nm in *B. subtilis* ^20,21^. The main component of the cell wall is the peptidoglycan, which consists of a polysaccharide backbone with β-(1,4) glycosidically linked N-acetylglucosamine (GlnNAc) and N-acetylmuramic acid (MurNAc) molecules ^22^. Attached to the lactyl group of N-acetylmuramic acid is an oligopeptide chain, which in most bacteria, including *Bacillus subtilis* and *Escherichia coli*, consists of L-Ala-D -Glu-L -meso-diaminopimelic acid-D -Ala-D-Ala ^23^.

Cell wall synthesis of the multilayered Gram positive cell wall is thought to occur at the cell membrane, with release of oldest strands to the cell periphery, and thus in an “inside-out” mode. In a first step, the soluble PG precursor consisting of a pentapeptide bound to UDP-MurNAc is synthesized in the cytoplasm. In the second phase, the linkage to undecaprenyl phosphate in the cytoplasmic membrane is catalysed, forming the lipid-linked monosaccharide peptide lipid I ^24^. Subsequently, the glycosyltransferase MurG ligates a N-acetyl-glucosamine (NAG) residue to lipid I generating the lipid-bound disaccharide-pentapeptide precursor, lipid II ^25^. In a next step, lipid II is transported across the plasma membrane to the outside by the flippase MurJ ^26^. In the final stage of cell wall biosynthesis, lipid II is polymerized and cross-linked by RodA and penicillin-binding proteins (PBPs) ^27^,^28^. In contrast to the Gram-negative cell wall, the Gram-positive cell wall possesses so-called WTAs (“wall teichoic acids”) and LTAs (“lipoteichoic acid”), that largely determine the charge of the cell wall. Their D-alanyl residues with free amino groups neutralize the negative charge of phosphates ^29^, making the cell wall more positively charged and influencing the folding and stability of secreted proteins by modulation of the availability of ions like *e*.*g*. calcium ^30^.

Visualization of protein secretion and the components of the secretion machinery has previously been studied up until the membrane barrier ^31^. We aim to advance the understanding of the location and dynamics of secretion including and focusing on cell wall passage. While we failed to visualize AmyE localization in the cell wall under native conditions, we were able to observe localized accumulation in the cell wall upon overproduction of AmyE, using two *Bacillus* species: *B. subtilis* and *B. licheniformis*. We argue that observed zones of protein secretion reflect genuine native regions of low density in the cell wall that allow for the diffusion of large proteins through the PG network.

## Results

### Secretion of active AmyE-mCherry in *B. subtilis* cells

It has been described that mCherry can be used as fluorescent reporter outside of the bacterial cell, *e*.*g*. within the periplasm of Gram negative bacteria ^32^. We sought to establish whether mCherry can be used as a reporter to visualize the secretion of proteins from *Bacillus* cells. We generated a fusion of AmyE-mCherry at the original gene locus, and observed only very weak fluorescence that could not be spatially resolved due to relatively high back ground fluorescence of *B. subtilis* in the red channel (data not shown). We resolved to using a high copy plasmid expressing AmyE-mCherry, which allowed yielding high concentrations of AmyE in the culture supernatant. We reasoned that as AmyE is heavily produced as well as secreted it might reveal its path across the cell envelope.

Fig. 1 shows that AmyE-mCherry when expressed from a plasmid, under the control of the strong constitutive HpaII promoter, is largely detected as full-length protein (72.6 + 26.7 kDa = 99.3 kDa) in *B. subtilis* cells, and to a large extent also in *B. licheniformis*, although here, degradation is quite extensive. As will become apparent below, expression at a single cell level is very heterogeneous, but expression levels of the entire population was quite similar between biological replicates (Fig. 1A). These experiments show that as long as AmyE-mCherry is associated with cells, it is protected from degradation by proteases present in the culture supernatant in *B. subtilis*, and can be efficiently detected in *B. licheniformis*.

**Fig. 1.**
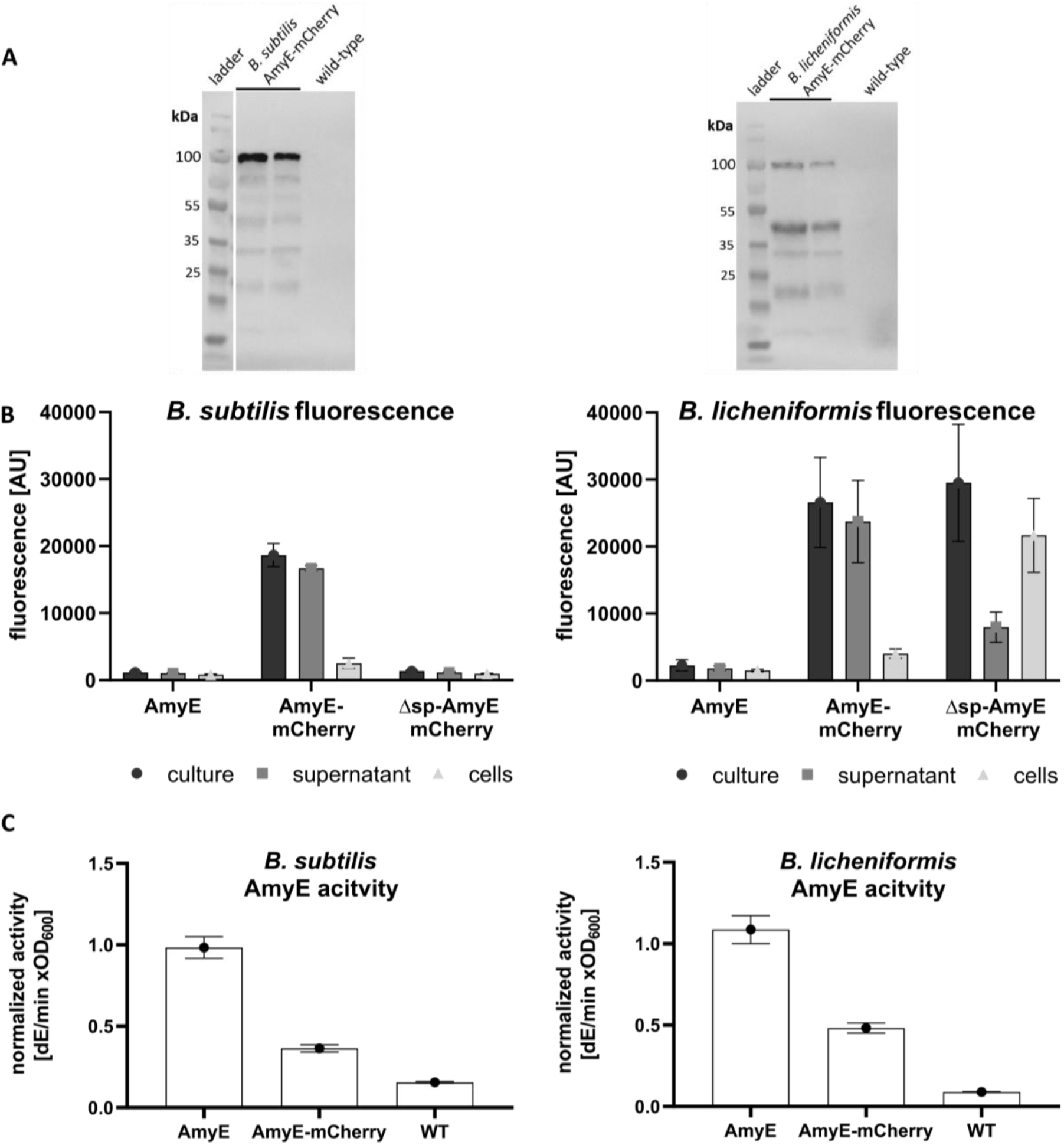
AmyE-mCherry is efficiently secreted from *B. subtilis* and *B. licheniformis* cells, but is also clearly detectable in a cell-associated from. (**A**) Western blot showing the presence of AmyE-mCherry fusion protein (calculated Mw: 99.3 kDa) in cell lysates of *B. subtilis* and *B. licheniformis* after 16 h of growth (note that duplicates are shown for assessment of reproducibility) using polyclonal antibodies against mCherry, (**B**) fluorescence measurement in whole culture, supernatant and separated cells, (**C**) amylase activity in culture supernatant. AmyE: plasmid-based expression of *amyE*, AmyE-mCherry: plasmid based expression of the reporter construct, WT: native genomic expression of *amyE* (C). Δsp-AmyE-mCherry represents a variant without signal peptide in (B).

In order to verify that AmyE-mCherry is secreted into the medium, we performed fluorescence assays of cultures grown to the transition to stationary phase (for a time course of secretion see Fig. 2B). Cells expressing AmyE from the native gene copy only showed weak background fluorescence, while cells expressing AmyE-mCherry from a plasmid displayed fluorescence. Strongest fluorescence was observed in the culture supernatant. When AmyE-mCherry was expressed as a polypeptide lacking the signal sequence, thus eliminating translocation across the cell membrane, fluorescence was reduced to background. In *B. licheniformis*, a very similar situation was observed. However, in the absence of a signal sequence clearly higher fluorescence was detected in the cells, but also in the culture supernatant, pointing to differences in the potential to fold and maintain the reporter protein in the cytoplasm between *B. subtilis* and *B. licheniformis*. (Fig. 1B). These findings suggest that while leaderless AmyE-mCherry accumulates within *B. licheniformis* cells, and is partially released, probably by cell lysis, secretion is lost in *B. subtilis*. Thus, fluorescence found in the supernatant of *Bacillus* cultures depends on the signal sequence of AmyE-mCherry, showing that the fusion protein is efficiently translocated across the cell membrane as well as across the cell wall. To prove that it is not only mCherry that is secreted, we determined amylase activity from culture supernatants. Higher amylase activity was determined for cells that produce AmyE from the plasmid than for wild type cells (Fig. 1C). Amylase activity was lower for cells overproducing AmyE-mCherry, indicating that the fusion either reduces AmyE activity, or reduced its efficiency of secretion. In any event, AmyE-mCherry activity was higher than in wild type cells, proving that AmyE-mCherry is efficiently secreted across the *B. subtilis* or *B. licheniformis* cell envelope and sufficiently stable for the following analysis.

**Fig. 2.**
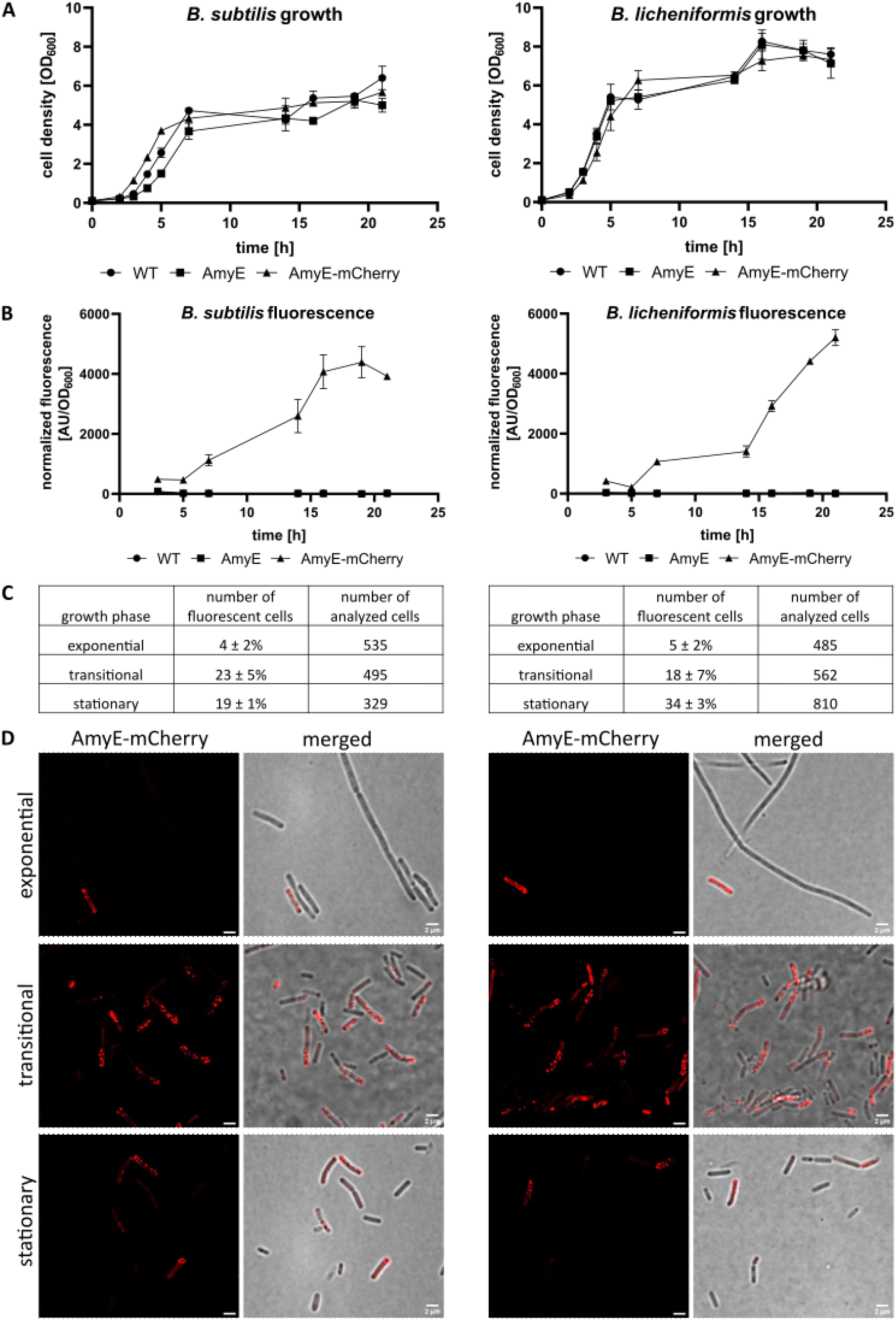
Dependence of cell density and fractions of cells producing AmyE-mCherry on growth phases. (**A**) Growth curves of *B. subtilis* and *B. licheniformis* incubated at 37°C for 21 h. (**B**) Kinetics of mCherry fluorescence of the whole culture (cell-associated plus culture supernatant) normalized to cell density. (**C**) Fraction of *B. subtilis* and *B. licheniformis* cells producing AmyE-mCherry in different growth phases. (**D**) Structured Illumination Microscopy (SIM) imaging of *B. subtilis* and *B. licheniformis* cells showing AmyE-mCherry fluorescence during different growth phases.

### AmyE-mCherry is secreted only by a subpopulation of cells

We next determined the expression profile of *Bacillus* cells overproducing AmyE-mCherry growing in liquid culture based on the fluorescence determined for the whole culture, *i*.*e*. cell-associated and in the culture supernatant. Fluorescence was observed in the culture starting with cells entering stationary phase, and continued to be well detectable in stationary growing cells (Fig. 2A). This is in line with reports of secretory processes usually commencing as cells transit from exponential into stationary phase ^33^.

We next sought to analyse the percentage of cells showing AmyE-mCherry fluorescence using fluorescence microscopy. During mid-exponential phase, we found 4 to 5% of cells showing AmyE-mCherry associated fluorescence in the red channel (Fig. 2B). This number increased during the transition phase to 23% and 18% for *B. subtilis* cells and *B. licheniformis* cells, respectively, (Fig. 2C). Interestingly, fluorescence was not found within the cytosol of cells, but was observed in a punctate pattern within the cell envelope, for both, *B. subtilis* as well as for *B. licheniformis*. These findings suggest that AmyE-mCherry accumulates at the cell membrane, and/or within the cell wall, but not within the cytosol.

During stationary phase, we observed an average of 34% of *B. licheniformis* cells showing AmyE-mCherry signals, while only 19% for *B. subtilis* cells (Fig. 2D).

Thus, high-level protein secretion is seen to occur in a highly heterogeneous manner, which is potentially also true for general protein secretion. We favour the idea that this is the case, because we find it unlikely that secretion of an overproduced protein should be restricted to a subpopulation of cells, while general protein secretion would operate in all cells.

### AmyE-mCherry is statically positioned in the cell envelope

We next investigated if the observed peripheral location of AmyE-mCherry foci is due to AmyE-mCherry being slowly translocated through the cell membrane, or to AmyE-mCherry being present within the cell wall. To address this question, we visualized the fusion protein in cells during transition phase, with or without treatment with lysozyme, which degrades to a large extent the *Bacillus* cell wall. Fig. 3 shows that more than 90% of cells treated with lysozyme lost rod shape and became sphaeroplasts upon treatment with lysozyme. While 23 ± 2% or 18 ± 6% of non-treated cells of *B. subtilis* and *B. licheniformis*, respectively, showed envelope-associated fluorescence of AmyE-mCherry, only 6 ± 3% or 3 ± 2% of cells, respectively, retained signals after exposure to lysozyme. We cannot distinguish if fluorescence seen in such sphaeroplasted cells is due to residual patches of peptidoglycan, or due to fusion proteins still attached to the cell membrane (via SecA and the translocon). Thus, envelope-associated signals of AmyE-mCherry are to a large extent due to the presence of intact PG layers, indicating that visible AmyE-mCherry accumulations at various positions along the cell wall represent protein molecules during the transit through the cell wall.

**Fig. 3.**
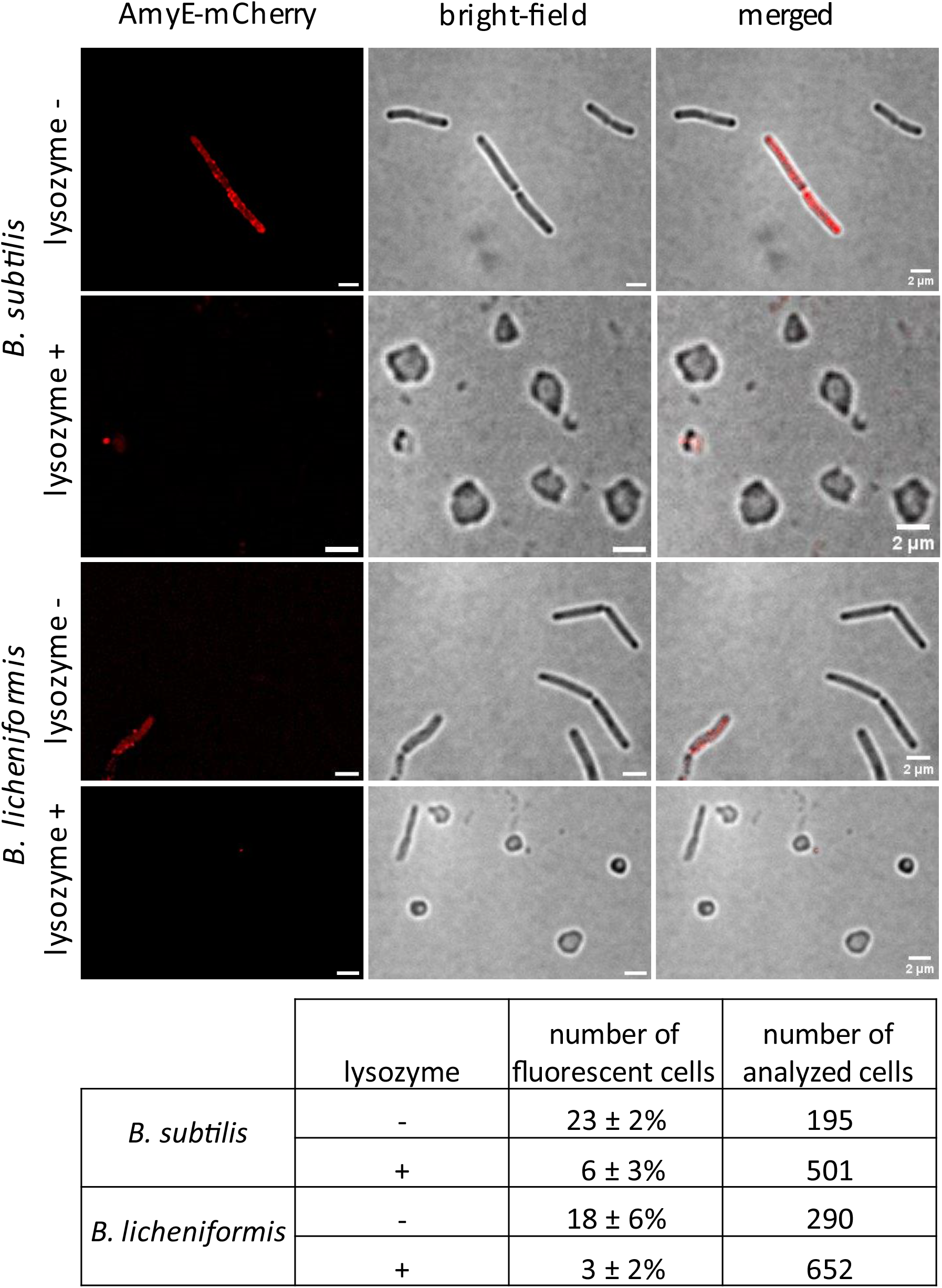
SIM imaging of AmyE-mCherry in *B. subtilis* and *B. licheniformis* in the transitional growth phase upon spheroplasting treatment with lysozyme. Cells displaying fluorescent signal were counted and normalized to the number of all analyzed cells.

### AmyE dynamics strongly differ from those of SecA or from SecDF

Intrigued by the observation that AmyE-mCherry fluorescence was recognized as punctate foci in two different *Bacillus* species, we set out to study the dynamics of the passage of the fusion protein through the cell wall. Because peptidoglycan synthesis is expected to occur from inside/out ^34-37^, and the wall to be rather rigid, we expected low lateral mobility of AmyE-mCherry foci. We employed structured illumination microscopy (SIM), reaching doubled resolution relative to conventional light microscopy. Time lapse experiments revealed that AmyE-mCherry foci remain statically positioned throughout many minutes (Fig. 4A). As opposed to this, even a slow-diffusing membrane protein (forming large clusters) such as flotillin FloT diffuses throughout the entire cell membrane of *B. subtilis* cells in time scale of 1.5 minutes ^38^. This finding suggests that AmyE-mCherry, after being transported across the cell membrane, does not equilibrate through the cell wall, even when produced in high amounts, but remains within distances of 125 nm or less. These data suggest that amylase secretion through the PG layers occurs within secretion zones, rather than throughout the cell wall. We tracked the position of foci relative to the long axis of cells using particle tracking. We found a speed of 0.97 pixels shift per frame (*i*.*e*. 60 s) for *B. licheniformis*, and a slightly lower speed of 0.87 for *B. subtilis*. To set this into relation, we tracked a SecA-mNeonGreen fusion expressed from the original gene locus in *B. subtilis* PY79 cells, as single copy of SecA in the cell. While *secA* is an essential gene in *B. subtilis*, cells grew indistinguishable from wild type cells, showing that the SecA-mNeonGreen fusion can functionally replace the wild type SecA copy. We also tracked a SecDF-NeonGreen fusion in the same genetic background. SecDF has been reported to play an important role in AmyE secretion, while not being essential for cell viability ^39^. Amylase activity in culture supernatant was similar in cells expressing SecDF-NeonGreen or the native copy of SecDF, indicating that the fusion was also largely functional (Fig. S1). SecA-NeonGreen formed discrete foci at the cell membrane in *B. subtilis* cells (a higher number than AmyE-mCherry), as has been described before for a SecA-GFP fusion ^31^, and these had a mean shift of 1.6 pixels/frame, almost twice as fast as AmyE-mCherry. SecDF-NeonGreen even showed a speed of two pixels per frame, more than two-fold higher than AmyE-mCherry. Thus, cytosolic and membrane proteins involved in AmyE secretion will come and go to and from AmyE secretion zones, *i*.*e*. to the SecYEG translocons involved in AmyE translocation across the cell membrane, while AmyE will continue to vertically diffuse through the wall towards the exterior of cells.

**Fig. 4.** AmyE-mCherry foci remain statically positioned. **A**) Time lapse SIM experiments of a *B. subtilis* cell expressing AmyE-mCherry. **B**) Single particle tracking of SIM time lapse images via TrackMate. SIM imaging for 10-30 minutes of AmyE-mCherry, SecDF-mNeonGreen and SecA-mNeonGreen fusion proteins in *B. subtilis* and AmyE-mCherry in *B. licheniformis*.

### AmyE secretion zones show fluctuations within a minutes-time scale

The cell wall has been estimated to have a thickness of 30 to 100 nm in Gram positive bacteria ^40^. The prevalence of many stationary AmyE-mCherry foci (remaining static for up to 30 minutes) suggests that AmyE-mCherry slowly diffuses through the lateral cell wall at several loci. When we analyzed time courses of AmyE-mCherry foci, we noticed that a considerable proportion of foci (25%) showed noticeable fluctuations in fluorescence intensity. Fig. 5 shows an example of *B. licheniformis* cells containing 2 foci that show strong fluctuations in fluorescence intensity. In order to rule out that fluctuations are caused by a shift in the focal plane or fluctuations in background fluorescence, we correlated fluorescence throughout cells with focal fluorescence intensity. Fig. 5B shows that fluctuations in focal fluorescence was largely independent of general fluctuations, and also much larger in intensity (as analyzed by converting into arbitrary units). Gain or loss of fluorescence could be observed between 1-minute intervals, suggesting that secretion zones can gain or lose fluorescent AmyE molecules within 60 second intervals. Fig. S2 shows more examples of such fluctuating structures within the *B. subtilis* cell wall. These data strongly support the notion of discrete zones within the cell wall that allow the passage of almost 100 kDa molecules within a time frame that is way below the scale of the 25 minutes for the cell cycle of cells growing in rich medium, not accounting for the fact that the cells are strongly slowing down growth during the transition into stationary phase. In this respect, a passive transport of secreted proteins through the meshwork of the murein sacculus by the incorporation of new peptidoglycan-precursors on the inner side of the cell wall and the gradual displacement of older glycan-strands to the outside as proposed by Kemper *et al*. ^41^ seems improbable

**Fig. 5.**
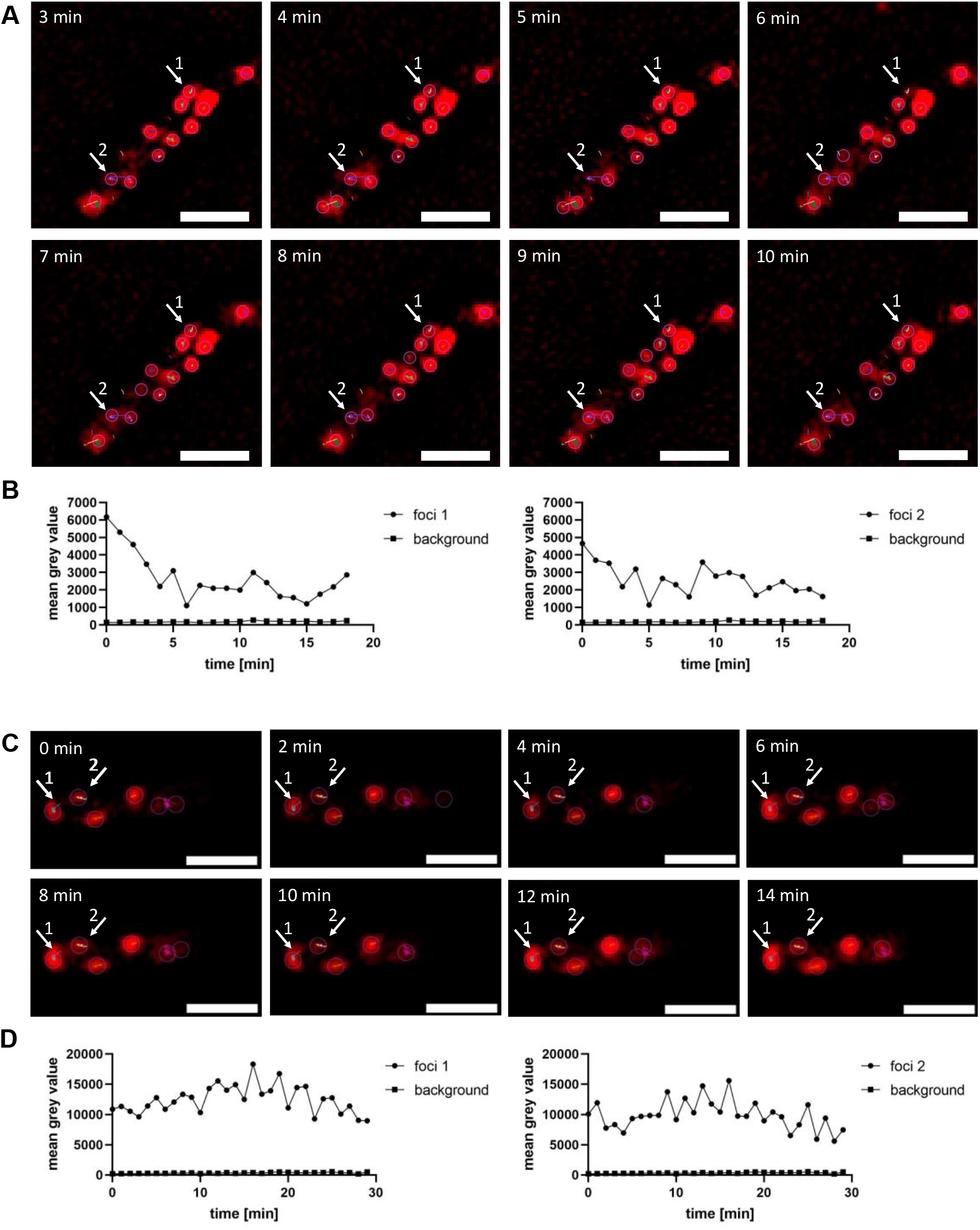
AmyE-mCherry foci showing intensity fluctuations over time. SIM time lapse images of AmyE-mCherry in (**A**) *B. licheniformis* and (**C**) in *B. subtilis*, showing each two foci that fluctuate in fluorescence intensity. (**B, C**) Fluorescence intensity analysis over time of the two foci and the background. Scalebar is 2 µm.

To test for the spatiotemporal involvement of SecA and SecDF in AmyE secretion, we analyzed the co-localization of SecA-mNeonGreen or SecDF-mNeonGreen fusions with AmyE-mCherry in cells during the transition phase. SecDF foci are found at almost all sites within the cell membrane (Fig. 6), thus might show a high degree of co-localization with AmyE secretion zones by default. However, using the program Fiji ^42^, we quantified 18 ± 3% spatial overlap between SecDF and AmyE, showing that AmyE-mCherry signals frequently lacked overlap with SecDF fluorescence. For SecA, the pattern of localization is more punctate. Fig. 6 shows that while SecA can be found in all cells, AmyE-mCherry foci cannot, in agreement with the heterogeneity of secretion zones observed (Fig. 2). This finding rules out that heterogeneous expression of SecA or of SecDF might be responsible for population-heterogeneity of secretion zones.

**Fig. 6.**
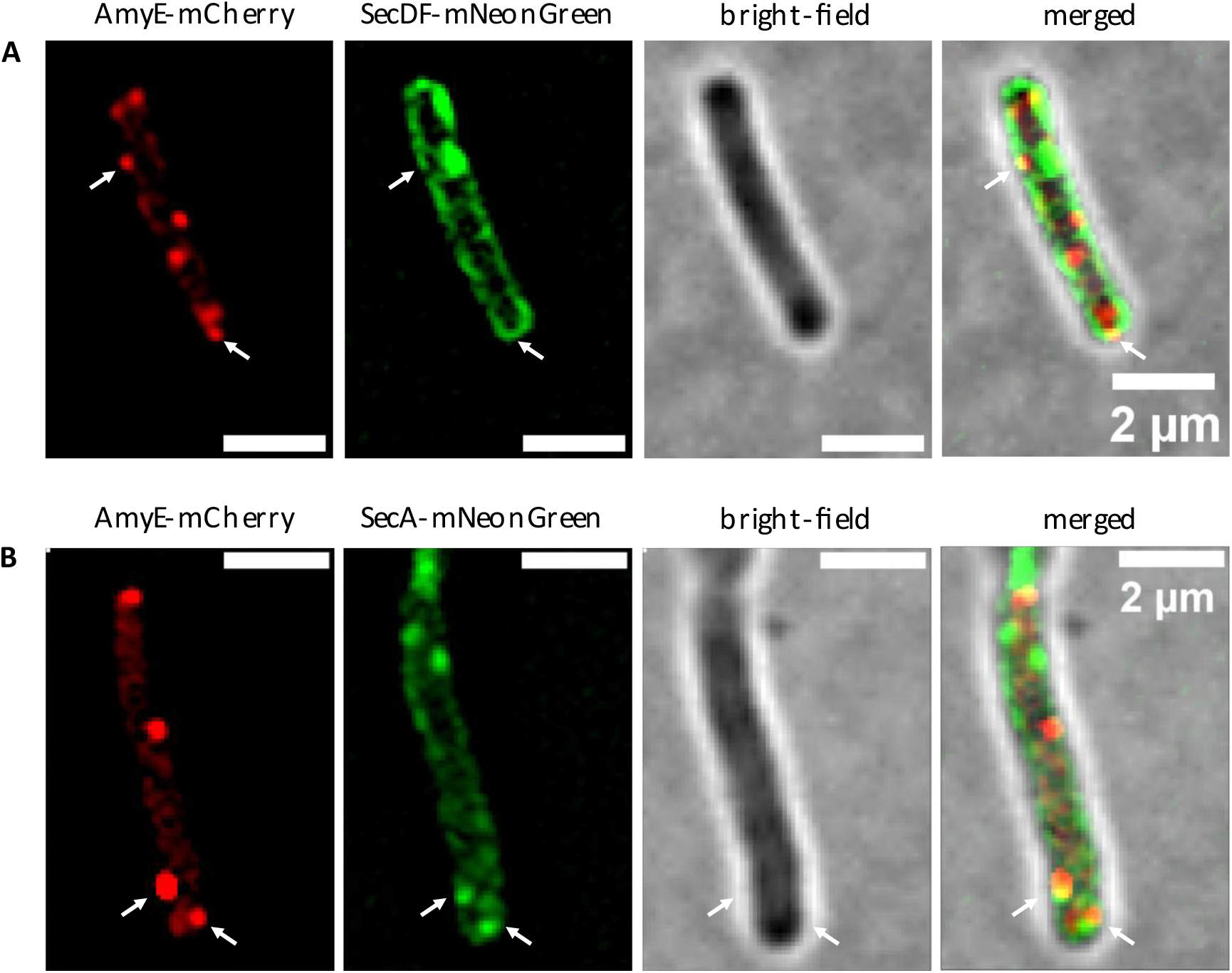
SIM imaging of *B. subtilis* cells co-expressing mCherry and mNeonGreen fusion. (**A**) Localization of AmyE-mCherry and SecDF-mNeonGreen with a shared area of 18% (3% SD) (**B**) Localization of AmyE-mCherry and SecA-mNeonGreen with a shared area of 19% (4% SD).

While there is co-localization of AmyE-mCherry and SecA-mNeonGreen foci, as expected from the essential nature of SecA within the secretion process, many AmyE-mCherry foci were observed lacking a SecA-mNeonGreen signal (Fig. 6). We quantified 19 ± 4% spatial overlap between both signals, revealing that secretion zones frequently lack an associated ATPase at the corresponding cytosolic site.

These observations are in agreement with the higher dynamics of SecA-mNeonGreen foci compared to AmyE-mCherry. SecA appears to accumulate at sites corresponding to AmyE secretion zones, but to diffuse to other sites once translocation across the SecYEG translocon has been achieved, or possibly even during the translocation of a single AmyE molecule. The data are also compatible with the idea that many AmyE molecules are translocated into the cell wall, where they diffuse towards the outside of the cell wall, constrained by the cell wall meshwork, and be released into the culture medium. Strong fluctuations of AmyE-mCherry fluorescence within secretion zones suggests that subsequently to loss of fluorescence via dissociation into the medium, new molecules can be translocated into a secretion zone, pointing to their long-lived nature, relative to assembly/disassembly of SecA foci at the cell membrane.

Our data support the hypothesis that AmyE molecules are guided into secretion zones and are released from them in a minutes’ time scale, as we corrected for a general loss or gain of fluorescence by fluctuations during image acquisition.

### AmyE is released from the cells at discrete zones

Our idea of secretion zones within the *Bacillus* cell wall implies that as amylase transits through the PG layers, it also emerges in defined zones from the cell envelope. To test this idea, we added fluorescently-labeled starch (“bodipy-starch”) to cells, an amylase substrate that develops fluorescence as it is degraded into monomers. Fig. 7 shows that fluorescence can be detected on 1% of *B. subtilis* cells carrying only the native copy of AmyE, nevertheless in a punctate manner. Overexpression of AmyE-mCherry gives rise to 18% and 25% of cells of *B. subtilis* and *B. licheniformis*, respectively, showing punctate fluorescent signals during the transition phase. Note that longer incubation with the substrate resulted in homogeneous staining of the cell surface (data not shown). These experiments indicate that as AmyE-mCherry enters secretions zones at the lower (inner) level of the cell wall, it exits the cell walls, as witnessed by its activity, at similarly discrete zones. Taken together, our data support the idea of areas of a diameter of 125 nm or less within the *Bacillus* cell wall, which allow the passage of many AmyE-mCherry molecules. We favour the view that this also holds true for the secretion of proteins produced at wild type-level, *i*.*e*. not overexpressed molecules.

**Fig. 7.**
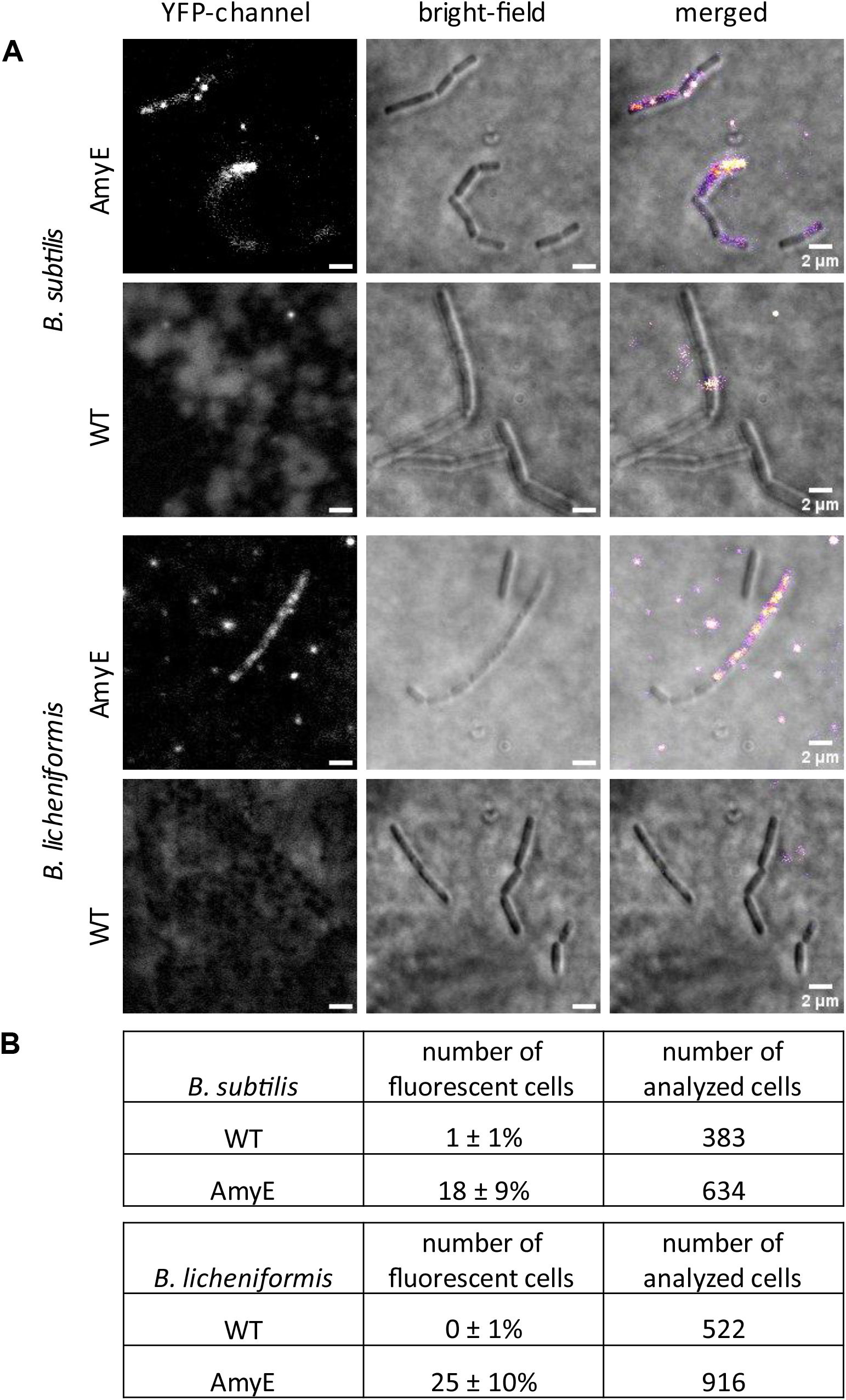
Localization of AmyE in *B. subtilis* and *B. licheniformis* cells determined by its activity. (**A)** Fluorescence is produced by hydrolysis of starch-BODIPY-FL when AmyE is secreted to the outer level of the cell wall. (**B**) Cells from three independent experiments displaying fluorescent signal were counted and normalized to the number of all analyzed cells.

### Expression of AmyE leads to considerable changes in SecA dynamics at a single molecule level

In order to better understand the SecA-driven process of AmyE secretion, we performed single molecule tracking (SMT) using the SecA-mNeonGreen fusion. SMT was performed as described before ^43,44^. SecA-mNeonGreen molecules were visualized using 20 ms stream acquisition, in cells grown to the transition phase (see movie S1). Using Squared Displacement Analyses (SQD) we found that the observed distribution of tracks was best fitted assuming three different populations of SecA-mNeonGreen molecules without overfitting of data (Fig. 8A, note that SMTracker 2.0 uses Bayesian Information Criterion and other tests to avoid overfitting artefacts). Fig. 8C displays the size of populations and their corresponding average diffusion constants from the data shown in Fig. 8A. Populations observed either moved with 0.7 µm^2^ s^−1^, a value compatible for a large, freely diffusing cytosolic protein (SecA forms a dimer in solution ^45,46^) ^47-49^, or with 0.15 µm^2^ s^−1^, in the range of freely diffusing ribosomal subunits ^50^, or with 0.04 µm^2^ s^−1^ (table 1). This extremely slow mobility has been proposed to account for the SRP system bound to the ribosome nascent chain complex as well as to the SecYEG translocon ^51^, or for translating ribosomes ^50,52^. According to this interpretation, about 21% of SecA is temporarily engaged in a secretion complex, while 50% diffuse through the cell and/or along the membrane bound to a substrate, and about 28% are freely diffusing, unbound SecA dimers. The three populations we observed are entirely compatible with data obtain by SMT on SecA from *E. coli* ^53^. During high AmyE secretion activity, SecA-mNeonGreen trajectories became shorter (Fig. 8A). The slow mobile “static” fraction of SecA-mNeonGreen increased from 20.9 to 29%, i.e. by 39%, while the medium “mobile” fraction remained relatively stable, and accordingly, the freely diffusing population decreased (Fig. 8C). These data suggest that most SecA molecules are bound to a substrate and in search of a translocon, and upon increased synthesis of AmyE, engagement with the translocon is increased, but free SecA is still available. In approximation to static engagement with a translocon, we determined average dwell time from the number of molecules staying within a radius of 106 nm (three times our localization error) for a given time. We determined about 300 ms for this time, no matter if AmyE was produced at wild type-level, or from the plasmid (table 1, Fig. 8G). Note that we are underestimating dwell times due to bleaching during imaging. The probability of dwell events could only be explained by using two exponential decay curves (Fig. 8G), suggesting that under wild type expression conditions, 78% of molecules have an average dwell time of 240 ms (*τ*_1_), and 22% of 450 ms (*τ*^2^). The latter fraction likely corresponds to molecules being bound to a translocon, in very good agreement with the population of static molecules (Fig. 8C, 21%); short dwell times can arise from stochastically occurring slow diffusion events. In cells carrying the plasmid overproducing AmyE, dwell times remained very similar (table 1), but the number of molecules showing extended dwell times (*τ*^2^, 430 ms) increased to 29%, again closely reflecting changes in population size of the static molecules (Fig. 8C). These finding suggest that while the time SecA spends on transport of molecules remains the same, more SecA molecules are engaged in transport events during AmyE overexpression.

**Table 1.** SMT data from SecA-mNeonGreen

|  | *B. subtilis* BG214 SecA-mNG | *B. subtilis* BG214 SecA-mNG AmyE |
| --- | --- | --- |
| Number of cells | 188 | 194 |
| average cell length [$\mu\text{m}$] | 2.7 | 2.8 |
| Number of *tracks* | 18455 | 11603 |
| population<sub>static</sub> [%] | $20.9 \pm 0.001$ | $29.0 \pm 0.001$ |
| population<sub>mobile</sub> [%] | $50.8 \pm 0$ | $50.4 \pm 0$ |
| population<sub>free</sub> [%] | $28.3 \pm 0.001$ | $20.6 \pm 0.001$ |
| D<sub>static</sub> [$\mu\text{m}^2 \cdot \text{s}^{-1}$] | $0.04 \pm 0$ | $0.04 \pm 0$ |
| D<sub>mobile</sub> [$\mu\text{m}^2 \cdot \text{s}^{-1}$] | $0.15 \pm 0$ | $0.15 \pm 0$ |
| D<sub>free</sub> [$\mu\text{m}^2 \cdot \text{s}^{-1}$] | $0.70 \pm 0.001$ | $0.70 \pm 0.001$ |
| R<sup>2</sup> | 1 | 1 |
| Average dwell time $\tau$ [s] | $0.297 \pm 0.004$ | $0.301 \pm 0.005$ |
| $\tau_1$ (2-comp.) [s] | $0.24 \pm 0.002$ | $0.23 \pm 0.002$ |
| fraction $\tau_1$ [%] | $78.2 \pm 1.95$ | $70.9 \pm 1.98$ |
| $\tau_2$ (2-comp.) [s] | $0.45 \pm 0.015$ | $0.43 \pm 0.012$ |
| fraction $\tau_2$ [%] | $21.8 \pm 1.95$ | $29.1 \pm 1.98$ |
|  | *B. subtilis* BG214 SecDF-mNG | *B. subtilis* BG214 SecDF-mNG AmyE |
| Number of cells | 161 | 83 |
| average cell length [$\mu\text{m}$] | 2,6 | 2,7 |
| Number of *tracks* | 11556 | 3566 |
| Population<sub>static</sub> [%] | $24,4 \pm 0,002$ | $29,7 \pm 0,002$ |
| Population<sub>mobile</sub> [%] | $46,9 \pm 0,001$ | $57,0 \pm 0,001$ |
| Population<sub>free</sub> [%] | $28,7 \pm 0,002$ | $13,0 \pm 0,002$ |
| $D_{\text{static}} [\mu\text{m}^2\cdot\text{s}^{-1}]$ | $0,03 \pm 0$ | $0,03 \pm 0$ |
| $D_{\text{mobile}} [\mu\text{m}^2\cdot\text{s}^{-1}]$ | $0,11 \pm 0$ | $0,11 \pm 0$ |
| $D_{\text{free}} [\mu\text{m}^2\cdot\text{s}^{-1}]$ | $0,60 \pm 0,002$ | $0,60 \pm 0,002$ |
| $R^2$ | 1 | 1 |
| Average dwell time $\tau$ [s] | $0,306 \pm 0,004$ | $0,288 \pm 0,005$ |
| $\tau_1$ (2-comp.) [s] | $0,25 \pm 0,001$ | $0,23 \pm 0,003$ |
| Fraction $\tau_1$ [%] | $80,7 \pm 1,36$ | $71,8 \pm 4,01$ |
| $\tau_2$ (2-komp.) [s] | $0,48 \pm 0,013$ | $0,39 \pm 0,017$ |
| Fraction $\tau_2$ [%] | $19,3 \pm 1,36$ | $28,2 \pm 4,01$ |

**Fig. 8.**
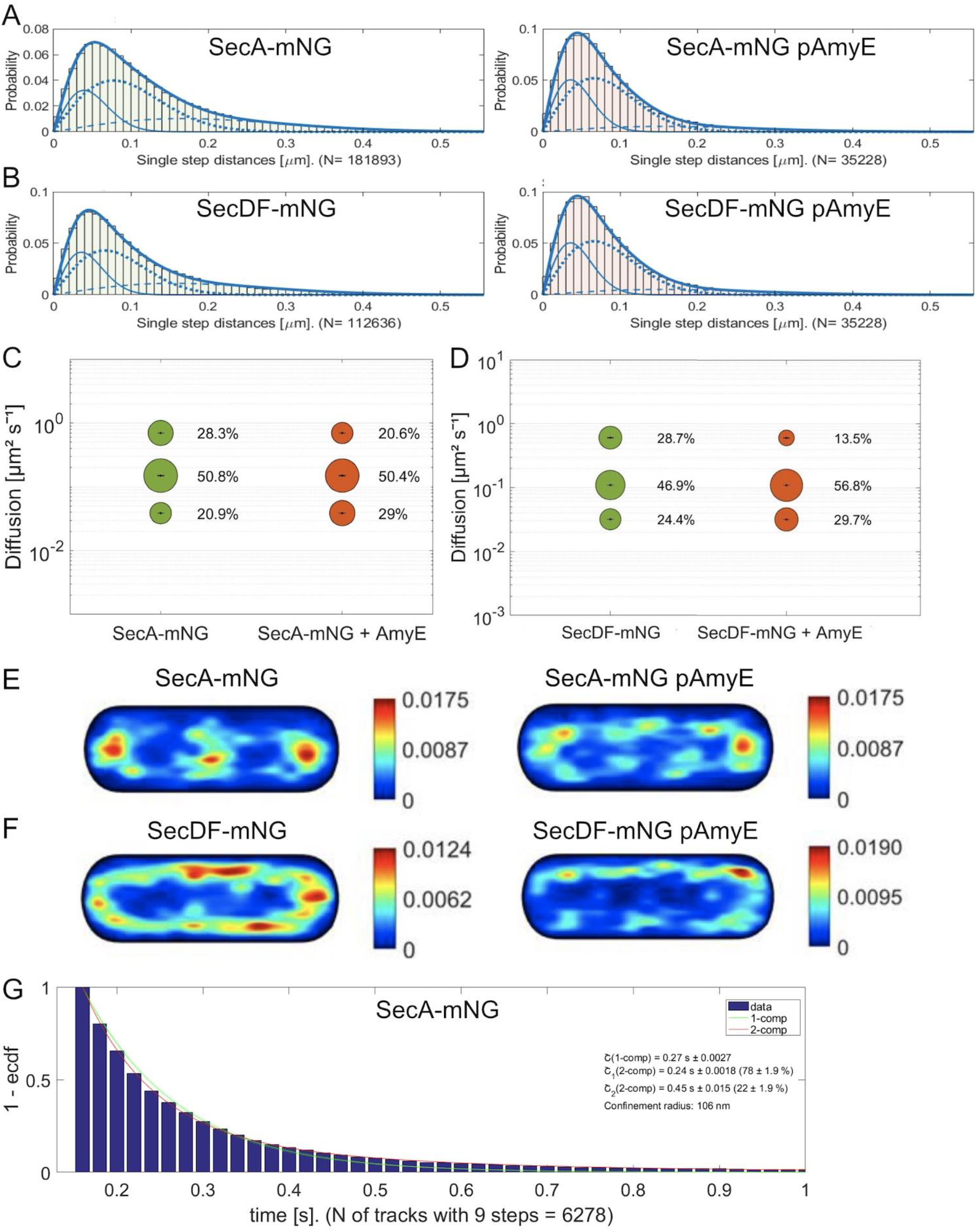
Single molecule tracking of SecA and SecDF. (A) Jump distance analyses of SecA-mNeonGreen (SecA-mNG) according to SQD analyses. Left panels represent wild type cells, right panels describe cells carrying the AmyE expression plasmid. This solid lines represent slow diffusing/static molecules, dotted lines are fits for medium-mobile molecules, dashed lines represent the fast-mobile population. (B) same as A for SecDF-mNeonGreen (mNG). (C) Bubble plots showing results from fitting of three populations by SQD analysis of single molecule tracks for SecA-mNeonGreen, size of bubbles correspond to population size, diffusion constants are given on the y-axis, bars in bubbles represent fitting errors. (D) bubble plot for SecDF-mNeonGreen data. (E) Heat map of static tracks of SecA-mNeonGreen projected into a 3 × 1 µm large cell, (F) similar as E) for SecDF-mNeonGreen. F) Plot of the probability density function of events of molecules staying within a radius of 106 nm for a certain amount of time (shown on the X-axis). The exponential decay curve can be explained by assuming two populations, one with a shorter (green curve) and one with a longer average dwell time (red curve), as stated in the inset.

When tracks were sorted into different populations, and tracks of the slow mobile “static” fraction were projected into a medium-sized cell of 3 × 1 µm, most molecules were found close to the cell poles, or in the cell centre, which is very similar to the localization of translating ribosomes ^50,54,55^ (Fig. 8C). Upon expression of plasmid-encoded AmyE, the pattern of localization of static SecA molecules changed, in that more sites along the lateral cell membrane showed high density of tracks (Fig. 8C).

SMT of SecDF also suggested the presence of three populations (Fig. 8B), of which the static population showed a milder increase upon overproduction of AmyE (Fig. 8D). The pattern of localization of static tracks became more uniform when cells expressed AmyE from plasmid (Fig. 8F, note the different scaling of the heat maps). Thus, SecDF also showed changes in single molecule dynamics during AmyE overproduction, but not as strongly as SecA.

These data support the idea that SecA exchanges between transport events at SecYEG translocons, in a time scale of few hundreds of milliseconds, in stark contrast to long-lived AmyE secretion zones, supporting the view that many AmyE molecules are transported into secretion zones by a highly dynamic population of SecA molecules.

## Discussion

We show that high-level secretion of amylase AmyE in two *Bacillus* species occurs at discrete zones throughout the cell envelope, mostly along the lateral side. Passage through the wall takes place at a minutes’ time scale, and appears to occur in a pulse-like manner. The finding of such protein secretion zones within the cell wall of low GC firmicutes has several far-reaching implications. Secretion zones require for locally large pores within the wall that extend perpendicular to the cell circumference allowing for the passage of 70 kDa (AmyE) and larger proteins. On average, the wall as a sieve-like meshwork of PG allows for the passage of up to 25 kDa proteins ^16^, and our data suggest that the wall is not homogeneously permissive for the passage of large proteins, but is discontinuous in its composition, including areas of lower meshwork density. Such structures have been hinted at by recent AFM visualization of the *B. subtilis* cell wall ^17,18^. Thus, a picture emerges that the multilayered PG envelope of firmicutes efficiently counteracts high intracellular turgor, but leaves many places for passage of proteins. We show that levels of AmyE-mCherry fluorescence change within a minute time frame, independent of fluorescence bleaching, showing decrease as well as increase. This finding suggests changes of numbers of amylase molecules within a secretion zone over time, consistent with constrained diffusion of a protein along a passage through a meshwork (of a thickness of about 50 nm) that slows down free diffusion through a solution, which occurs in a time frame of milliseconds for nanometer distances ^56^. As such, secretion zones appear to allow for faster diffusion as opposed to the typically pictured homogeneous PG meshwork.

A caveat in our analyses is that AmyE-mCherry signals could only be discerned from back ground fluorescence that is relatively strong in *B. subtilis* cells in the red fluorescence channel (stronger than *e*.*g*. in *E. coli* cells), *i*.*e*. during overproduction of AmyE-mCherry from a high copy number plasmid. It could be argued that accumulation of AmyE within discrete zones in the cell wall could be an artefact of protein overproduction. We suggest that this is not the case, based on the following considerations: we observed secretion zones at the transition to stationary phase, when *B. subtilis* is known to secrete a multitude of proteins, from proteases to lipases, including several sugar-polymer-degrading enzymes ^57^. It is unlikely that overexpression of one of these proteins disturbs or overwhelms the entire system. Also, we did not observe noticeable changes in growth of cells that could point to a stress situation. At the time of visual identification of secretion zones, the transition to stationary growth, cell wall synthesis has stopped, or is at least strongly slowing down ^37^. It is hard to envision that cell wall structure should strongly change during a phase of arrest of synthesis. We propose that increased synthesis of AmyE allows us to track the path of molecules, as opposed to low production level, which does not allow tracking the passage of fewer molecules versus back ground fluorescence. Fluctuations of AmyE-mCherry fluorescence also suggests that secretion zones are not clogged up with overproduced AmyE molecules, but allow for an oscillating passage of many molecules, including bursts of release and phases of re-accumulation, through gaps in the PG structure.

Heterogeneity of transcriptional expression of genes is a well-established phenomenon in bacteria ^58^, as well as share of labour between cells growing in a biofilm ^59^. While some cells provide energy to generate extracellular matrix in biofilms, others engage in competence development or spore formation, or remain mobile and ensure dispersing of cells from biofilms ^60^. Production of antibiotics has been shown to occur in a heterogeneous manner ^61^, and even DNA repair enzymes can be found in only a subset of exponentially growing cells, leading to heterogeneity of DNA damage response, in this case based on extremely low numbers of molecules per cell ^62^. Likewise, c-di-GMP signaling components of *B. subtilis* cells are found to be absent in a considerable subpopulation of cells, due to low abundance of proteins within the network ^63^. The mentioned phenomena of heterogeneity notwithstanding, we were surprised to see that overproduction of AmyE-mCherry follows a very strong pattern of heterogeneity, with a maximum of 23% of cells showing AmyE-mCherry secretion zones during the transition phase, and 34% during stationary phase. Heterogeneity was observed as cells entered stationary growth, but was not based on heterogeneity of SecA expression in cells. We cannot exclude the possibility that plasmid loss in a fraction of cells contributes to this heterogeneity. For the plasmid-based production of proteins in *B. megaterium* fluctuating plasmid abundance was observed, which resulted in population heterogeneity ^64^. However, as plasmid loss requires cell division events, and heterogeneity in case of AmyE arose as cells turned off cell division, plasmid loss is unlikely to contribute to a major degree.

In addition to larger cavities within the cell wall observed from isolated cell walls 18, secretion zones may increase in size and number as cells turn off cell wall synthesis, in a heterogeneous manner. Zones containing larger pore sizes of the peptidoglycan meshwork may put cells at risk of bursting due to internal turgor. A culture entering stationary phase may thus be evolved to allow for a minority of cells going at risk of dying, in order to provide large amounts of extracellular enzymes for the rest of the population.

AmyE secretion zones did not show lateral mobility within the cell, in agreement with the presence of structures within the cell wall that allow for molecule passage. While SecA also showed the formation of focal assemblies at the cell membrane, these showed higher lateral dynamics than AmyE secretion zones, and likewise, SecDF showed much higher dynamics at the cell membrane. These data suggest that SecA molecules move between SecYEG translocons (for which we have so far failed to generate functional fusions), transporting AmyE molecules, which carry on moving through the wall at these sites. In order to obtain a better spatiotemporal resolution of SecA dynamics, we employed single molecule tracking. As was described for *E. coli* SecA ^53^, we found three populations of SecA molecules having strongly different average diffusion constants. These populations can be best explained by molecules actively transporting secreted proteins at the translocon (about 20%), SecA molecules having bound cargo in search of a translocon (about 50%), and freely diffusing SecA dimers (30%). Upon overproduction of AmyE, the slow mobile population increased to about 30%, the medium mobile fractions remained constant, and the freely diffusing molecule decreased to 20%, suggesting that more SecA molecules are engaged in active transport, but that there is still a substantial pool of free SecA molecules to allow for efficient general protein secretion to continue. Interestingly, average dwell times of SecA did not change, but the population of about 20% of molecules showing a longer dwell time increased to about 30% upon AmyE overproduction, suggesting that average transport times remain constant (as well as exchange of SecA molecules between translocons), but the number of molecules dwelling at the translocon increases.

*Overall*, our data strongly support the findings of cell wall heterogeneity, show that a subpopulation of cells secretes amylase at discrete zones, allowing proteins to move through the wall, in a manner compatible with Brownian motion. This would also explain why a putative machinery allowing active or directed transport of proteins through the *Bacillus* cell wall has never been identified. In contrast to slow AmyE dynamics, SecA shows high turnover at SecYEG translocons, and becomes more engaged during AmyE overproduction, but is not overwhelmed with additional AmyE secretion. Thus, protein secretion in *Bacilli* is a two-tier process involving very different time scales of protein motion.

## Material & Methods

### Bacterial strains and plasmids

*The B. subtilis* strain used was PY79 (derivative of Bacillus 168), the *B. licheniformis* MC28 and MC26 strains were provided by B.R.A.I.N. Biotech AG (Zwingenberg, Germany) (Supplementary Table S1). MC26 was used as a control strain. Bacillus strains were grown at 37°C overnight on nutrient agar plates using commercial nutrient broth LB solidified by addition of 1% (w/v) agar. Overnight cultures in tubes were inoculated from a fresh agar plate and incubated overnight at 37°C and 200 rpm. Day cultures in 100 ml shake flasks with 10 ml media were inoculated to a cell density of OD_600_ of 0.1 in LB from the overnight cultures and then incubated at 37°C and 200 rpm.

For the visualization of the secreted protein α-amylase AmyE, the mCherry gene was cloned via Gibson Assembly in frame to *amyE* in plasmid pM11K_amyEBs provided by the B.R.A.I.N. AG (Zwingenberg, Germany). This plasmid provides the HpaII-promoter ^65^ to drive the expression of *amyE* and a high copy number pUB110-like replicon. The C-terminal fusion includes an 8-amino acid linker (KLGSGSGS). This non-integrating plasmid carries a kanamycin resistance for selection with 25 µg/ml kanamycin in *Bacillus*. The plasmid is available, upon reasonable request, after signing a Material Transfer Agreement.

The fusion of SecA and SecDF to mNeonGreen were cloned into the pSG1164 vector containing a sequence encoding monomeric NeonGreen ^66^ and a flexible 14-amino acid linker (GPGLSGLGGGGGSL). For this purpose, at least 500 bp of the 3‵ end of the desired gene (excluding the stop codon) was amplified by polymerase-chain reaction (PCR) using *B. subtilis* PY79 gDNA as template, oligonucleotides (Supplementary Table S2), Phusion DNA polymerase, and deoxynucleotide solution (both from New England Biolabs, NEB). The resulting PCR product was integrated into the plasmid via the Gibson Assembly cloning system (New England Biolabs-NEB). The pSG1164 plasmid integrates at the native locus of the corresponding gene by a single-crossover event, creating a C-terminal fusion ^67^.

### Structured Illumination Microscopy (SIM)

Samples taken typically at transitional growth phase were mounted on ultrapure-agarose slides dissolved in LB (1%) for immobilization of cells prior to image acquisition. For localization experiments, image Z-stacks (∼100 nm steps) were acquired using brightfield (BF) image acquisition (transmitted light) or structured illumination microscopy (SIM) with a ZEISS ELYRA PS.1 setup (Andor EMCCD camera, 160 nm pixel size; 3× rotations and 5× phases per z-slice; with an excitation wavelength 561 nm at 15% intensity or 488 nm at 10% intensity; ZEISS alpha Plan-Apochromat 100x/NA 1.46 Oil DIC M27 objective). SIM reconstructions were processed using ZEN-Black software by ZEISS. ImageJ2/FIJI version 1.52p was used for visualization and image processing ^42,68 69^. For time lapse imaging the acquisition time was set to 1 minute. SIM reconstructions were manually cropped in axial and lateral dimensions, depending on plausibility of cellular positions, using “Duplicate”-function. Signal not connected to the cells was considered to be background and was therefore in most cases eliminated. For single-particle tracking, spots were identified with the LoG Detector of TrackMate v6.0.1 [Tinevez et al., 2017], implemented in Fiji 1.53 q, an estimated diameter of 0.5 μm and sub-pixel localization activated. Spots were merged into tracks via the Simple LAP Tracker of TrackMate, with a maximum linking distance of 500 nm, one frame gaps allowed, and a gap closing max distance of 800 nm.

### Generation of protoplasts

*Bacillus* cells in the transitional growth phase were treated according to the protocol of Chang & Cohen ^70^ to obtain protoplasts. During the process kanamycin was added to the media to maintain the AmyE-mCherry plasmid. Imaging of the cells before and after the incubation with lysozyme was performed by SIM microscopy.

### Starch-BODIPY-FL staining

For this experiment the streptococcal SpeB protocol for *Streptococcus* by Rosch & Caparon ^71^ was adapted to *Bacillus*. Strains were cultivated in LB medium at 37°C and 200 rpm with the addition of 25 µg/ml kanamycin until the transitional growth phase. The culture was pelleted at 4000 rpm for 2 min and resuspended in fresh LB containing 1% of the “DQ starch substrate stock solution” (1 mg/ml, EnzChek Ultra Amylase Assay Kit, Invitrogen Detection Technologies, Carlsbad, CA, USA). Cells were mounted on ultrapure-agarose slides dissolved in LB (1%) for immobilization of cells and incubated for 30 minutes at 37°C.

Imaging was performed via epi-fluorescence microscopy, using a Nikon Eclipse Ti-E, Nikon Instruments Inc with a CFI Apochromat objective (TIRF 100× oil, NA 1.49) and an EMCCD camera (ImagEM X2 EM-CCD, Hamamatsu Photonics KK). The samples were illuminated with Nikon C-HGFIE Intensilight (Precentered Fiber Illuminator) and the YFP-channel filter cube ET 500/20, T 515 LP, ET 535/30. Images were processed with MetaMorph (version 2.76), and ImageJ ^42^.

### Phadebas test for amylase activity

For the quantification of α-amylase activity in the culture supernatant, the Phadebas Amylase Test (Phadebas AB, Uppsala, Sweden) was used. One Phadebas tablet was dissolved in 20 ml buffer solution (0.1 M acetic acid, 0.1 M potassium acetate, 5 mM calcium chloride, pH 5). Overnight cultures of *Bacillus* were centrifuged at 14000 rpm for 2 minutes in a microfuge, 20 µl supernatant was mixed with 180 µl substrate solution and incubated for 10 min at 37°C and 1000 rpm in a thermomixer (Eppendorf Thermomixer comfort). The reaction was stopped by addition of 60 µl 1 M sodium hydroxide. The reaction tubes were centrifuged and the absorption of 100 µl of the supernatant measured at 620 nm via a microplate reader (Tecan Infinite 200 PRO, Tecan, Switzerland). Activities were corrected for dilution and normalized to the cell density (OD_600_) of the culture.

### Immunoblotting

30 ml of a culture in the transitional growth phase was pelleted and resuspended in 3 ml buffer (100 mM NaCl, 50 mM EDTA, 5 mg/ml Lysozyme). Cells were incubated at 37°C until lysis, which was observed visually. Samples were incubated at 95°C with sodium dodecyl sulfate (SDS) loading buffer for 5 minutes. Proteins were separated by polyacrylamide gel electrophoresis (PAGE) on a 10% mini-PROTEAN^®^ TGX^™^ precast gel (Bio-Rad, CA, USA) at 140 V and 300 mA for 1 h. Gels were transferred onto cellulose membranes using transfer-buffer (48 mM Tris, 39 mM glycine, 1.3 mM SDS, 20% EtOH, pH 9.8) at 25 V, 500 mA for 1 h. Membranes were blocked for 1 h using blocking-buffer (PBS, 0.1% Tween-20 with 5% w/v nonfat dry milk) and incubated with diluted (1:10.000) rabbit polyclonal antiserum (Sigma-Aldrich) against mCherry overnight. Subsequently, membranes were washed three times with PBS for 5 minutes each and incubated with goat-anti-Rabbit-IgG, peroxidase-conjugated (1:10.000) for 1 h (Sigma-Aldrich). Before detection of proteins the membranes were washed three times as described before. Detection was performed using an Immobilon^®^ Forte Western membrane substrate (Merck KGA, Darmstadt, Germany) according to the manufacturer’s protocol. Protein marker Thermo Scientific™ PageRuler Prestained Protein Ladder was used.

### Single molecule tracking

Individual molecules were tracked using custom-made slim-field setup on an inverted fluorescence microscope (Nikon Eclipse Ti-E, Nikon Instruments Inc.). An EMCCD camera (ImagEM X2 EM-CCD, Hamamatsu Photonics KK) was used to ensure high-resolution detection of the emission signal, resulting in a calculated resolution of the position of the molecule down to 20 nm. The central part of a 514 nm laser diode (max power 100 mW, TOPTICA Beam Smart) was used with up to 20% of the intensity (about 160 W cm-^2^ in the image plane) to excite samples, fused to mNeonGreen (using laser filter set BrightLine 500/24, dichroic mirror 520 and BrightLine 542/27), by focusing the beam onto the back focal plane of the objective. A CFI Apochromat objective (TIRF 100 x Oil, NA 1.49) was used in the setup ^72^. For the analysis, a video of 3000 frames at 20 ms was recorded, of which the last 1000 frames were used for the analysis. Software Oufti ^73^ was used to set the necessary cell meshes. Utrack ^74^ was employed for automatic determination of molecule trajectories. Data analysis was carried out using software SMTracker 2.0 ^72,75^.

## Acknowledgments

We gratefully acknowledge help with SIM microscopy by Sven Holtrup and Maximilian Greger from SYNMIKRO, Marburg University. This work was supported by the Bundesministerium für Bildung und Forschung (BMBF, Program NatLifE).

## Materials & Correspondence

Correspondence and material requests should be addressed to Peter Graumann.

## Author contributions

MS performed all experiments, except for those shown in Fig. 8; FK and SF performed single molecule tracking experiments shown in Fig. 8; MS, KL and PLG wrote the manuscript; KL and PLG conceived of the study and obtained funding for the work. PLG supervised experiments.

## Data availability

All data are shown the manuscript. Raw single molecule data are available as http://dx.doi.org/10.17192/fdr/111.

